# Visual features as stepping stones toward semantics: Explaining object similarity in IT and perception with non-negative least squares

**DOI:** 10.1101/029314

**Authors:** Kamila M. Jozwik, Nikolaus Kriegeskorte, Marieke Mur

**Author notes:** Email addresses or.

## Abstract

Object similarity, in brain representations and conscious perception, must reflect a combination of the visual appearance of the objects on the one hand and the categories the objects belong to on the other. Indeed, visual object features and category membership have each been shown to contribute to the object representation in human inferior temporal (IT) cortex, as well as to object-similarity judgments. However, the explanatory power of features and categories has not been directly compared. Here, we investigate whether the IT object representation and similarity judgments are best explained by a categorical or a feature-based model. We use rich models (> 100 dimensions) generated by human observers for a set of 96 real-world object images. The categorical model consists of a hierarchically nested set of category labels (such as “human”, “mammal”, “animal”). The feature model includes both object parts (such as “eye”, “tail”, “handle”) and other descriptive features (such as “circular”, “green”, “stubbly”). We used nonnegative least squares to fit the models to the brain representations (estimated from functional magnetic resonance imaging data) and to similarity judgments. Model performance was estimated on held-out images not used in fitting. Both models explained significant variance in IT and the amounts explained were not significantly different. The combined model did not explain significant additional IT variance, suggesting that it is the shared model variance (features correlated with categories, categories correlated with features) that best explains IT. The similarity judgments were almost fully explained by the categorical model, which explained significantly more variance than the feature-based model. The combined model did not explain significant additional variance in the similarity judgments. Our findings suggest that IT uses features that help to distinguish categories as stepping stones toward a semantic representation. Similarity judgments contain additional categorical variance that is not explained by visual features, reflecting a higher-level more purely semantic representation.

## 1. Introduction

Inferior temporal (IT) neurons in primates are thought to respond to visual image features of intermediate complexity, consisting of object parts, shape, color, and texture (Komatsu et al. 1992; Kobatake & Tanaka, 1994; Tanaka 1996; Kayaert et al. 2003; Yamane et al. 2008; Freiwald et al. 2009; Issa & DiCarlo 2012). Consistent with this selectivity profile, moderately scrambled object images activate human IT almost as strongly as their intact counterparts (Grill-Spector et al. 1998). These findings suggest that object representations in IT are feature-based. However, the literature on human IT (Kanwisher et al. 1997; Epstein et al. 1998; Haxby et al. 2001; Downing et al. 2001; Kriegeskorte et al. 2008b; Mur et al. 2012) has stressed the importance of category membership in explaining IT responses. Object category membership is a characteristic of the whole object, and requires a representation that is invariant to variations in visual appearance among members of the same category. Many studies have indicated that category membership of perceived objects can explain a significant proportion of the IT response variance, at the level of single neurons (e.g. Tsao et al. 2006), and, more strongly, at the level of brain regions (e.g. Kanwisher et al. 1997; Epstein et al. 1998; Tsao et al. 2003; Mur et al. 2012) and neuronal population codes (e.g. Haxby et al. 2001; Hung et al. 2005; Kiani et al. 2007; Kriegeskorte et al. 2008b).

The representationin a neuronal population code can be characterized by its representational geometry (Kriegeskorte et al. 2008a; Kriegeskorte & Kievit 2013). The population’s representational geometry is defined by the distance matrix among the representational patterns and reflects what stimulus properties are emphasized and de-emphasized in the representation. The IT representational geometry has been shown to emphasize certain category divisions that are behaviorally relevant to a wide variety of species, including the division between animate and inanimate objects and, within that, between faces and bodies (Kriegeskorte et al. 2008b; Kiani et al. 2007). Additional support for the importance of categories in shaping IT comes from the fact that successful modeling of IT responses to natural objects appears to require a categorical component of one form or another. Until recently, models using categorical labels (provided by humans) clearly outperformed image-computable models in predicting IT responses (e.g. Naselaris et al. 2009; Huth et al. 2012). Recently, deep convolutional neural networks trained on category-discrimination tasks to achieve high performance (e.g. Krishevzky et al. 2012) have been shown to explain the IT representation better than any previous image-computable models (Kriegeskorte, in press; Yamins et al. 2014; Khaligh-Razavi & Kriegeskorte 2014; Cadieu et al. 2014).

Despite the importance of both features and categories in the human and primate IT literature, there is little work directly comparing the explanatory power of features and categories for explaining the IT representation. Given that the categorical structure in the IT object representation must emerge from constituent object parts and features, both types of information may account for variance in the IT representational geometry. Recent observations have indeed also suggested the existence of a continuous component in the IT object representation (Kriegeskorte et al. 2008b; Connolly et al. 2012; Mur et al. 2013; Sha et al. 2015). The continuous component might for instance be driven by object shape variations (Op de Beeck et al. 2001; Haushofer et al. 2008; Drucker & Aguirre 2009). The presence of a continuous component hints at an underlying feature-based code. The idea that feature-based population coding might underlie a categorical representation is consistent with previous cognitive theory and experimental work (Tyler & Moss, 2001; Op de Beeck et al. 2008a; Tsunoda et al. 2001; Vogels et al. 1999), and with the proposal that IT contains feature detectors optimized for category discrimination (Sigala & Logothetis 2002; Ullman et al. 2002; Ullman 2007; Lerner et al. 2008).

A second related question is which type of representation best explainsperceived object similarity. Perceived object similarity has been shown to reflect both the continuous and categorical components of the IT object representation (Edelman et al. 1998; Op de Beeck et al. 2001, 2008b; Haushofer et al. 2008; Mur et al. 2013). However, this leaves open what the relative contributions of visual features and categories are to perceived object similarity. Possible clues come from classic psychophysics work, which suggests an important role for category information in object perception (e.g. Rosch et al. 1976). Moreover, object similarity judgments are more strongly categorical than the IT object representation and show additional category divisions not present in the IT representation, including the division between human and nonhuman animals, and between manmade and natural objects (Mur et al. 2013).

Here we investigate the extent to which features and categories or a combination of both can account for object representations in IT and for object similarity judgments. We constructed a feature-based and a categorical model from object descriptions generated by human observers for a set of 96 real-world object images (the same set as used in Kriegeskorte et al. 2008b). The categorical model consists of a hierarchically nested set of category labels (such as “human”, “mammal”, “animal”). The feature model includes both object parts (such as “eye”, “tail”, “handle”) and other descriptive features (such as “circular”, “green”, “stubbly”). These rich models (114 category dimensions, 120 feature-based dimensions) were fitted to the brain representation of the objects in IT and early visual cortex (based on functional magnetic resonance imaging data), and to human similarity judgments for the same set of objects. The models were fitted using non-negative least squares and tested on independent sets of images. Figure 1 shows a schematic overview of model creation and fitting. We used representational similarity analysis (Kriegeskorte et al. 2008a; Nili et al. 2014) to compare the performance of the feature-based and categorical models in explaining the IT representation and the similarity judgments.

**Figure 1.**
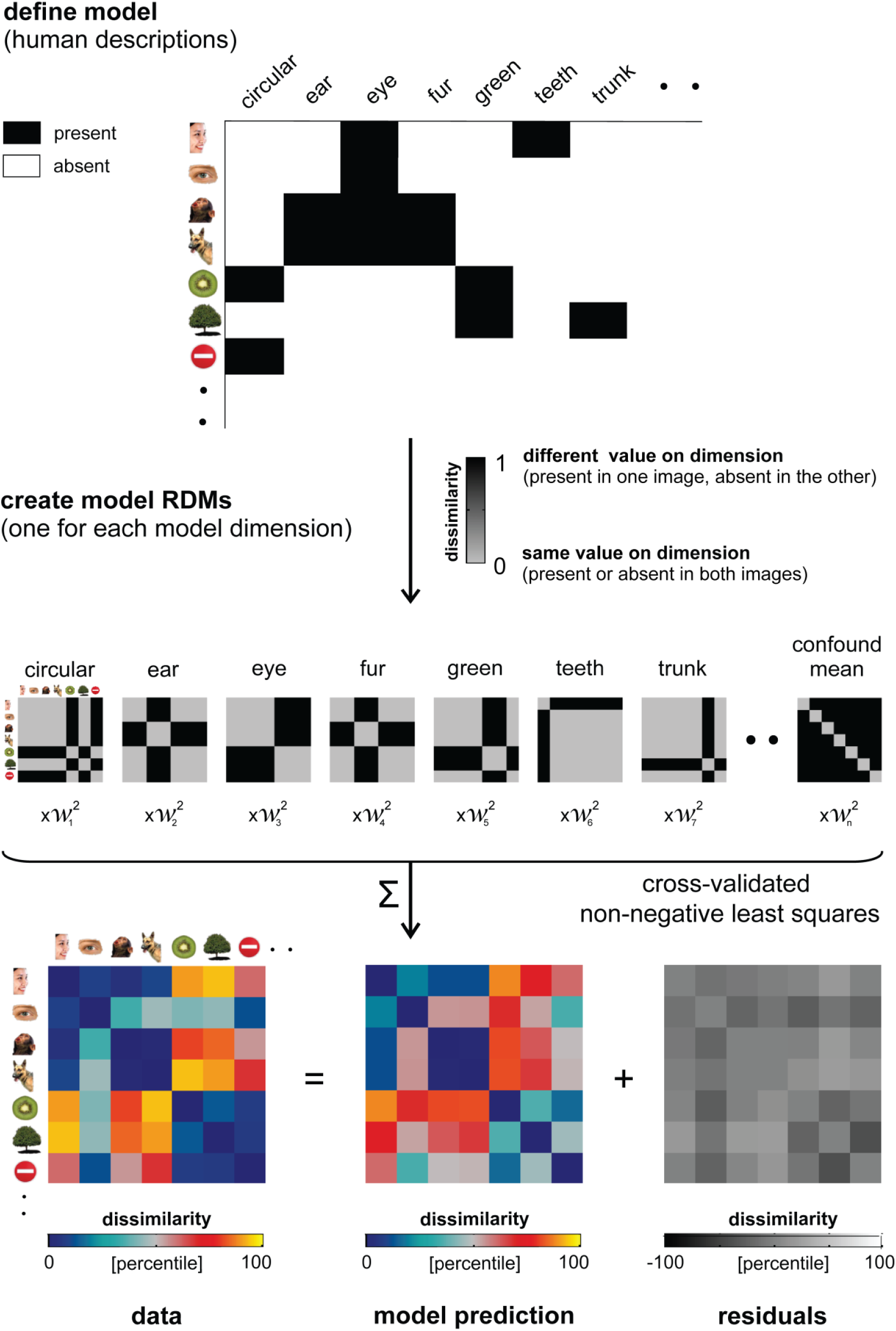
Schematic overview of model creation and fitting. The schematic shows a set of example images and feature-based model dimensions. We defined the model dimensions (e.g. “circular”, “ear”), and the value of each image on these dimensions, by asking human observers to generate and verify image descriptions. We subsequently created a model RDM for each dimension, which indicates for each pair of images whether they have the same or a different value on that dimension. Finally, we implemented non-negative least squares (LS) fitting to find the single-dimension model-RDM weights that optimally predict the data RDM. Each model includes a confound-mean predictor. The weights were estimated using a cross-validation procedure to prevent over fitting.

## 2. Methods

### 2.1 fMRI experiment

Acquisition and analysis of the fMRI data have been described in Kriegeskorte et al. (2008b), where further details can be found.

#### 2.1.1 Subjects

Four healthy human volunteers participated in the fMRI experiment (mean age = 35 years; two females). Subjects were right-handed and had normal or corrected-to-normal vision. Before scanning, the subjects received information about the procedure of the experiment and gave their written informed consent for participating. The experiment was conducted in accordance with the Institutional Review Board of the National Institutes of Mental Health, Bethesda, MD.

#### 2.1.2 Stimuli

Stimuli were 96 colored images of objects from a wide range of categories, including faces, animals, fruits, natural scenes, and manmade objects. The stimuli are shown in Supplementary Figure 1.

#### 2.1.3 Experimental design and task

Stimuli were presented using a rapid event-related design (stimulus duration, 300 ms; inter stimulus interval, 3700 ms) while subjects performed a fixation-cross-color detection task. Stimuli were displayed on a uniform gray background at a width of 2.9° visual angle. Each of the 96 object images was presented once per run. Subjects participated in two sessions of six nine-minute runs each. In addition, subjects participated in a separate block-localizer experiment. Stimuli (grayscale photos of faces, objects, and places) were presented in 30-s category blocks (stimulus duration, 700 ms; interstimulus interval 300 ms). Subjects performed a one-back repetition-detection task on the images.

#### 2.1.4 Functional magnetic resonance imaging

Blood-oxygen-level-dependent fMRI measurements were performed at high resolution (voxel volume: 1.95×1.95×2 mm^3^), using a 3 Tesla General Electric HDx MRI scanner, and a custom-made 16-channel head coil (Nova Medical Inc.). We acquired 25 axial slices that covered inferior temporal (IT) and early visual cortex bilaterally (single-shot, gradient-recalled Echo Planar Imaging: matrix size: 128 × 96, TR: 2s, TE: 30ms, 272 volumes per run, SENSE acquisition).

#### 2.1.5 Estimation of single-image activity patterns

fMRI data were preprocessed in BrainVoyager QX (Brain Innovation) using slice-scan-time correction and head-motion correction. All further analyses were conducted in Matlab (The MathWorks Inc.). Single-image activity patterns were estimated for each session by voxel-wise univariate linear modeling (using all runs except those used for region-of-interest definition). The model included a hemodynamic-response predictor for each of the 96 stimuli along with run-specific motion, trend and confound-mean predictors. For each stimulus, we converted the response-amplitude (beta) estimate map into a t map.

#### 2.1.6 Definition of regions of interest

All regions of interest (ROIs) were defined on the basis of independent experimental data and restricted to a cortex mask manually drawn on each subject’s fMRI slices. IT was defined by selecting the 316 most visually-responsive voxels within the inferior temporal portion of the cortex mask. Visual responsiveness was assessed using the t map for the average response to the 96 object images. The t map was computed on the basis of one third of the runs of the main experiment within each session. To define early visual cortex (EVC), we selected the 1057 most visually responsive voxels, as for IT, but within a manually defined anatomical region around the calcarine sulcus within the cortex mask. EVC does not show a clear categorical structure in its responses, and was therefore included in our analyses as control region.

#### 2.1.7 Construction of the representational dissimilarity matrix

For each ROI, we extracted a multivoxel pattern of activity (t map) for each of the 96 stimuli. For each pair of stimuli, activity-pattern dissimilarity was measured as 1 minus the Pearson linear correlation across voxels within the ROI (0 for perfect correlation, 1 for no correlation, 2 for perfect anticorrelation). The resulting 4560 pairwise dissimilarity estimates were placed in a representational dissimilarity matrix(RDM). RDMs were constructed for each subject and session separately and then combined by averaging across sessions and subjects. The RDMs capture the information represented by a brain region by characterizingits representational geometry (Kriegeskorte et al. 2008a; Kriegeskorte & Kievit, 2013). The representational geometry of a brain region reflects which stimulus information is emphasized and which is deemphasized.

### 2.2 Object-similarity judgments

Acquisition and analysis of the object-similarity judgments have been described in Mur et al. (2013), where further details can be found.

#### 2.2.1 Subjects

Sixteen healthy human volunteers participated in the similarity-judgment experiment (mean age = 28 years; 12 females). Subjects had normal or corrected-to-normal vision; 13 of them were right-handed. Before participating, the subjects received information about the procedure of the experiment and gave their written informed consent for participating. The experiment was conducted in accordance with the Ethics Committee of the Faculty of Psychology and Neuroscience, Maastricht University, The Netherlands.

#### 2.2.2 Stimuli

Stimuli were the same 96 object images as used in the fMRI experiment. The stimuli are shown in Supplementary Figure 1.

#### 2.2.3 Experimental design and task

We acquired pair wise object-similarity judgments for the 96 object images by asking subjects to perform a multi-arrangement task (Kriegeskorte & Mur, 2012). During this task, the object images are shown on a computer screen in a circular arena, and subjects are asked to arrange the objects by their similarity, such that similar objects are placed close together and dissimilar objects are placed further apart. The multi-arrangement method uses an adaptive trial design, showing all 96 object images on the first trial, and selecting subsets of objects with weak dissimilarity evidence for subsequent trials. In other words, the method will “zoom in” to objects that were placed close together on previous trials. The multi-arrangement method allows efficient acquisition of a large number of pair wise similarities. Each subject performed the task for one hour. In the instruction, we intentionally did not specify which object properties to focus on, as this would have biased our perspective on the mental representation of the objects.

#### 2.2.4 Construction of the representational dissimilarity matrix

Subjects were instructed to use the entire arena on each trial. Consequently, only the relations between distances on a single trial, not the absolute on-screen distances, were meaningful. For each subject, dissimilarity estimates were therefore averaged across trials using an iterative procedure, alternately scaling the single-trial estimates to match their evidence-weighted average, and recomputing the evidence-weighted average, until convergence (Kriegeskorte & Mur, 2012).RDMs were constructed for each subject separately and then combined by averaging across subjects. The resulting RDM captures which stimulus information is emphasized and which is de-emphasized in object perception.

### 2.3 Defining the categorical and feature-based models

We performed two behavioural experiments to obtain the categorical and feature-based models. In Experiment 1, a group of human observers generated category and feature descriptions for the 96 object images. These descriptions are the model dimensions. In Experiment 2, a separate group of human observers judged the applicability of each model dimension to each image, thereby validating the dimensions generated in Experiment 1, and providing, for each image, its value (present or absent) on each of the dimensions. The images’ values on the validated model dimensions define the model. Figures 2 and 3 show the categorical and feature-based models, respectively.

**Figure 2.**
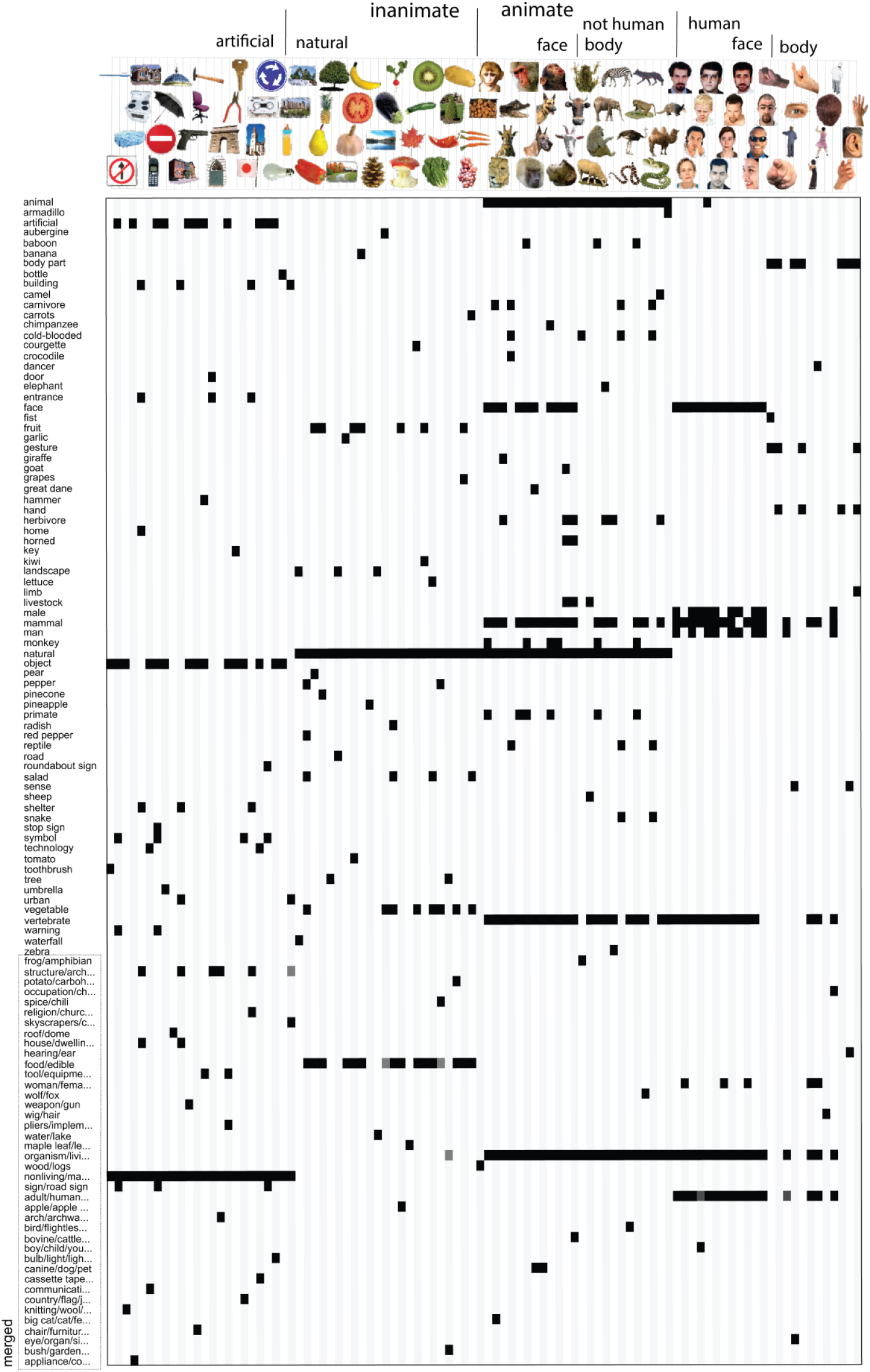
Categorical model. Rows correspond to model dimensions (114 in total); columns correspond to the 96 object images. Each image is centered with respect to the column that it corresponds to (e.g. the first column shows the values of the tooth brush on each dimension). Black indicates that a category is present; white indicates that it is absent. Gray values might appear for merged dimensions. For display purposes, the labels of some of the merged dimensions are truncated. The labels are listed in full in Appendix E.

#### 2.3.1 Experiment 1: Object descriptions

Fifteen healthy human volunteers participated in Experiment 1 (mean age = 26 years; 11 females). Subjects were native English speakers, right-handed, and had normal or corrected-to-normal vision. Before participating, the subjects received information about the procedure of the experiment and gave their written informed consent for participating. The experiment was conducted in accordance with the Cambridge Psychology Research Ethics Committee, Cambridge, United Kingdom.

During the experiment, we asked subjects to generate descriptions, of categories and features, for the 96 object images. In the instruction, we defined a category as “a group of objects that the shown object is an example of”. The instructions further stated that an object can belong to multiple categories at once, with categories ranging from specific to more and more abstract. We defined features as “visible elements of the shown object, including object parts, object shape, color and texture”. The instruction contained two example images, not part of the 96 object-image set, with category and feature descriptions. We asked subjects to list a minimum of five descriptions, both for categories and for features. See Appendix A for detailed subject instructions.

The entire measurement session took three hours, approximately equally divided between the generation of category and feature descriptions. The order of the two tasks was counterbalanced across subjects. The 96 images were shown, in random order, on a computer screen using a web-based implementation, with text boxes next to each image for subjects to type category or feature descriptions. Subjects could scroll down to move to the next few images, and press a button when they were done to save their data.

We subsequently selected, for categories and features separately, those descriptions that were generated by at least three out of 15 subjects. This threshold corresponds to the number of subjects that on average mentioned a particular category or feature for a particular image. The threshold is relatively lenient, but it allows inclusion of a rich set of descriptions, which were further pruned in Experiment 2.Wesubsequently removed descriptions that were either inconsistent with the instructions or redundant. After this step, there were 197 category descriptions and 212 feature descriptions. These descriptions are listed in Appendix B and C.

#### 2.3.2 Experiment 2: Validation

Fourteen healthy human volunteers participated in Experiment 2 (mean age = 28 years; seven females). Subjects were native English speakers and had normal or corrected-to-normal vision. Thirteen of them were right-handed. Before participating, the subjects received information about the procedure of the experiment and gave their written informed consent for participating. The experiment was conducted in accordance with the Cambridge Psychology Research Ethics Committee, Cambridge, United Kingdom.

The purpose of Experiment 2 was to validate the descriptions generated during Experiment 1. We therefore asked an independent group of subjects to judge which descriptions correctly described which images. During the experiment, the object images and the descriptions, each in random order, were shown on a computer screen using a web-based implementation. The object images formed a column, while the descriptions formed a row; together they defined a matrix with one entry, or checkbox, for each possible image-description pair. We asked the subjects to judge for each description, whether it correctly described each object image, and if so, to tick the associated checkbox. Subject could scroll up and down and left to right while they were going through the images and descriptions, and press a button when they were done to save their data. See Appendix D for detailed subject instructions.

A measurement session took approximately three hours, during which a subject would only have time to judge either the category or the feature descriptions. Of the 14 subjects, six judged category descriptions, six judged feature descriptions, and the remaining two judged both. This resulted in eight subjects for the category validation experiment and eight subjects for the feature validation experiment.

We subsequently kept, for categories and features separately, those image-description pairs that were judged as correct by at least six out of eight subjects. This relatively strict threshold aims at including only those image descriptions that can generally be expected to be judged as correct. This procedure creates a binary vector for each description with length equal to the number of object images, where 1 indicates that the description applies to the image (present), and 0 indicates that it does not (absent). Descriptions whose resulting binary vectors only contained zeros (i.e. they were not ticked for any image by at least six people) were removed. This reduced the number of descriptions to 179 for the categories, and 152 for the features. We subsequently removed any obvious incorrect ticks, which mainly involved category-related ticks during the feature validation experiment (e.g. ticking “hammer” for an image of a hammer instead of for an image of a gun). As a final step, to increase the stability of the weights estimated during regression, we iteratively merged binary vectors that were highly correlated (r >0.9), alternately computing pair wise correlations between the vectors, and averaging highly-correlated vector pairs, until all pair wise correlations were below threshold. The resulting set of 114 category vectors forms the categorical model (Figure 2) and the resulting set of 120 feature-based vectors forms the feature-based model (Figure 3). Merged vectors might contain values in the range (0 1). The final sets of descriptions are listed in full in Appendix E.

**Figure 3.**
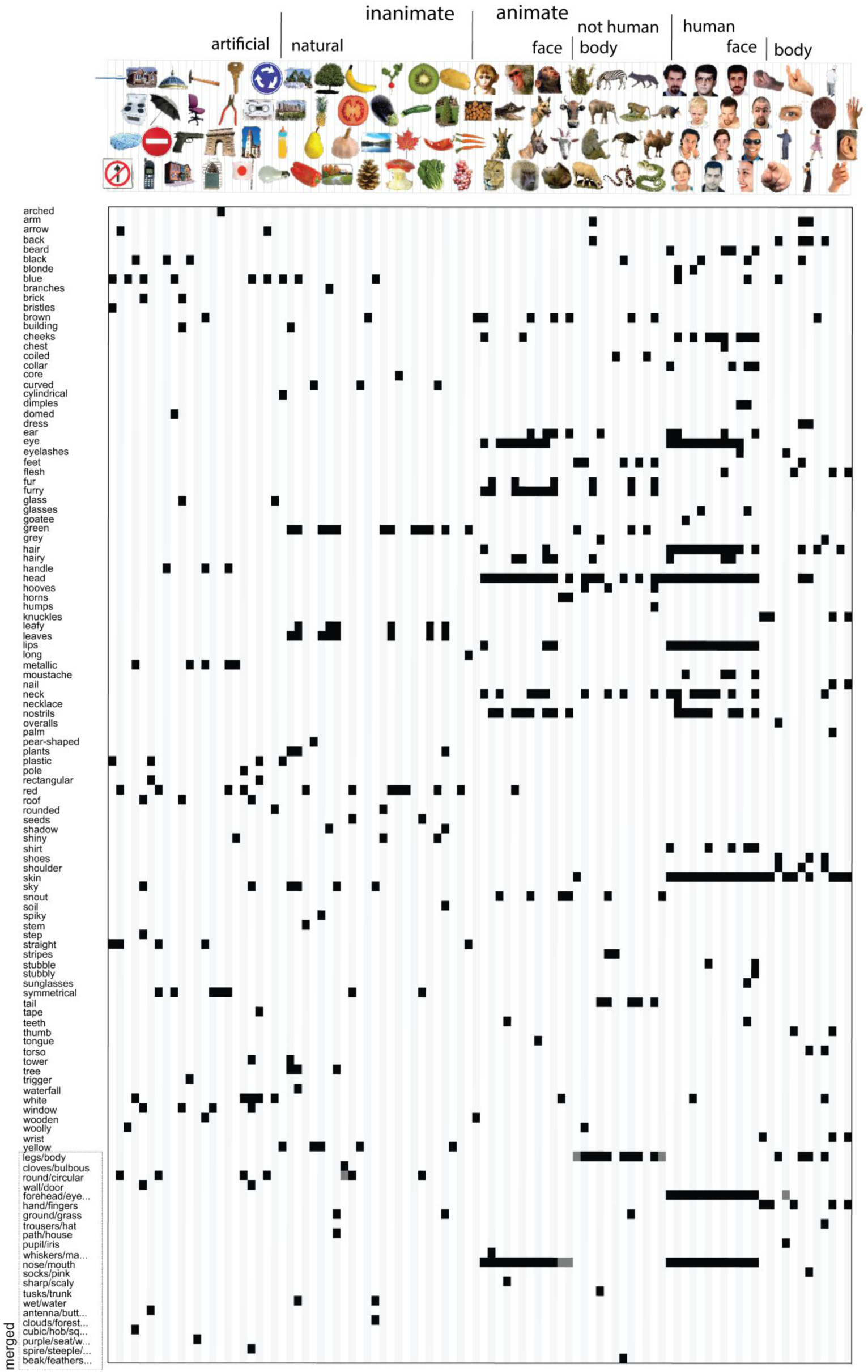
Feature-based model. Rows correspond to model dimensions (120 in total); columns correspond to the 96 object images. Each image is centered with respect to the column that it corresponds to (e.g. the first column shows the values of the toothbrush on each dimension). Black indicates that a feature is present; white indicates that it is absent. Gray values might appear for the merged dimensions. For display purposes, the labels of some of the merged dimensions are truncated. The labels are listed in full in Appendix E.

#### 2.3.3 Creatingmodel RDMs

In order to compare the models to the measured brain representation and similarity judgments, the models and the data should reside in the same representational space. This motivates transforming our models to “RDM space”: for each model dimension, we computed, for each pair of images, the absolute difference between their values on that dimension. The absolute difference reflects the dissimilarity between the two images in a pair. Given that our models are practically binary, almost all dissimilarities were either 0 or 1. A dissimilarity of 0 indicates that two images have the same value on a dimension, i.e. the category or feature is present or absent in both images. A dissimilarity of 1 indicates that two images have a different value on a dimension, i.e. the category or feature is present in one image, and absent in the other. Figures 4 and 5 show the single-dimension model RDMs for the categorical and feature-based model, respectively.

**Figure 4.**
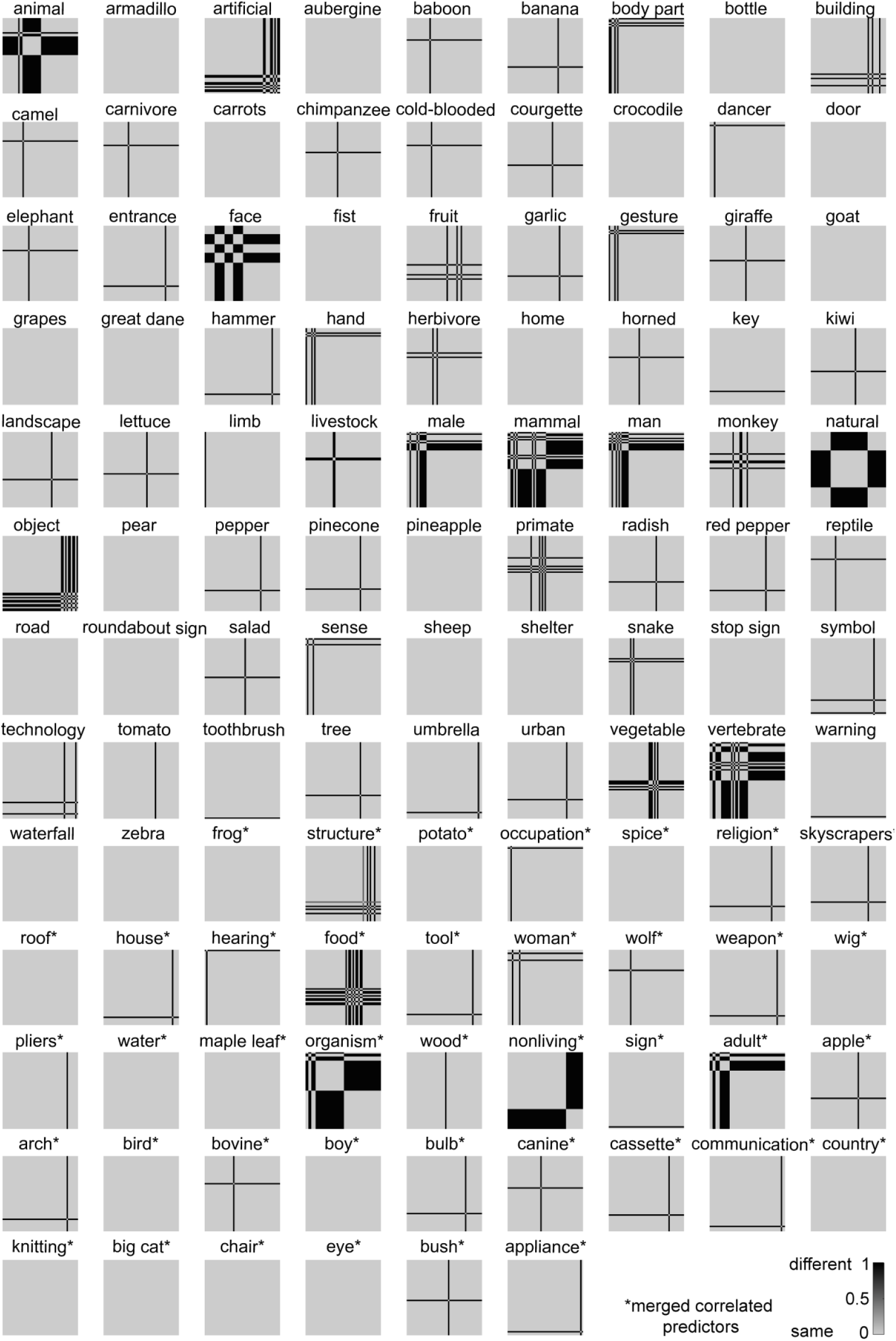
Single-dimension model RDMs of the categorical model. The single-dimension model RDMs were created by determining for each dimension (i.e. each column in Figure 2) which object pairs have the same value (category present or absent for both objects; dissimilarity = 0) and which object pairs have a different value (category present for one object, and absent for the other; dissimilarity = 1). Dissimilarity values in the range (0 1) might appear for merged dimensions. For merged dimensions only the first label of the merged set is shown. Merged dimensions are indicated with an asterisk.

**Figure 5.**
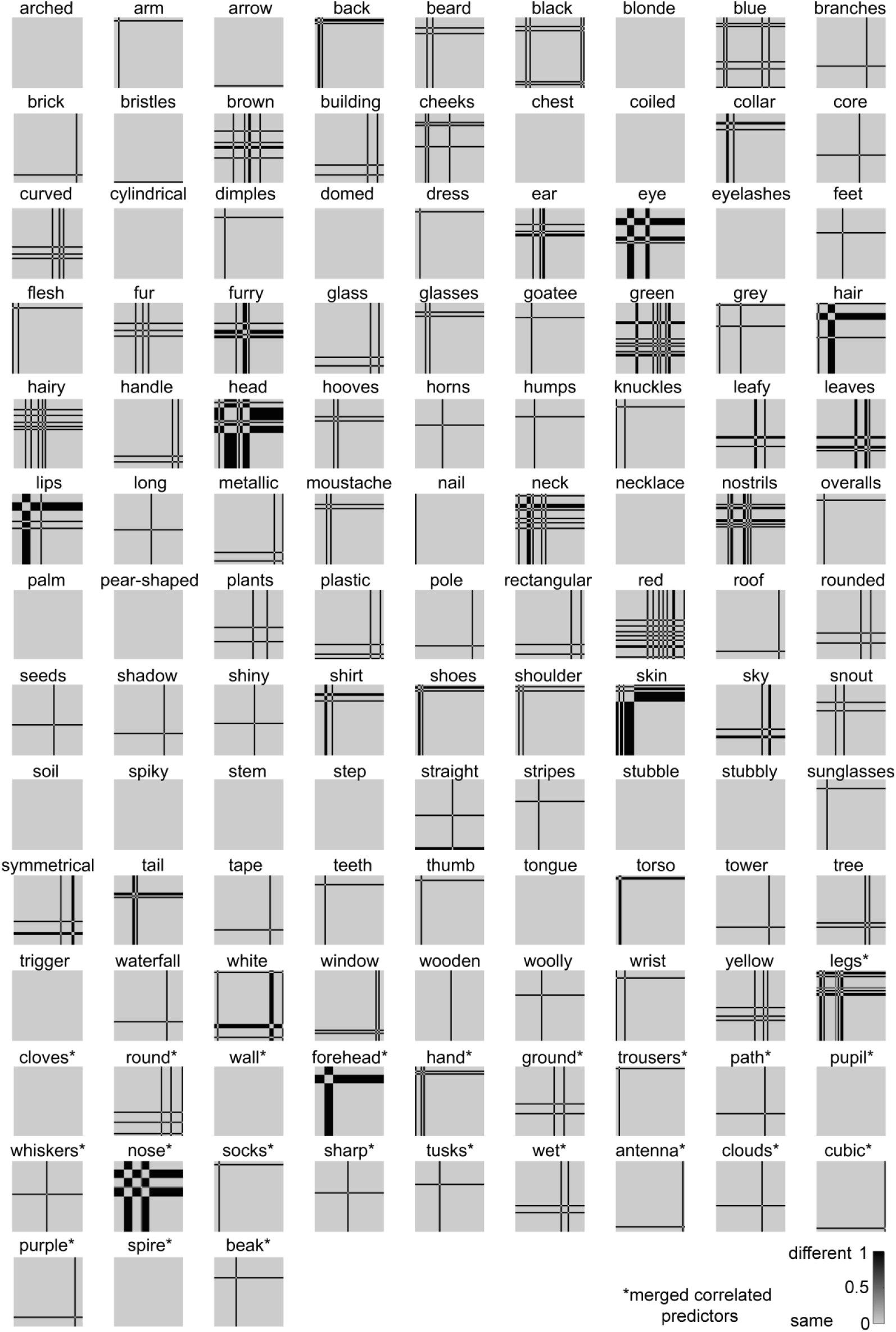
Single-dimension model RDMs of the feature-based model. The single-dimension model RDMs were created by determining for each dimension (i.e. each column in Figure 3) which object pairs have the same value (feature present or absent for both objects; dissimilarity = 0) and which object pairs have a different value (feature present for one object, and absent for the other; dissimilarity = 1). Dissimilarity values in the range (0 1) might appear for merged dimensions. For merged dimensions only the first label of the merged set is shown. Merged dimensions are indicated with an asterisk.

### 2.4 Non-negative least-squares fitting of the representational models

We could predict the brain representation and dissimilarity judgments by making the assumption that each model dimension contributes equally to the representation. We use the squared Euclidean distance as our representational dissimilarity measure, which is the sum across dimensions of the squared response difference for a given pair of stimuli. The squared differences simply sum across dimensions, so the model prediction would be the sum of the single-dimension model RDMs. However, we expect that not all model dimensions contribute equally to the brain representation or similarity judgments. This motivates weighting the model dimensions to optimally predict the measured object representation. This approach not only increases the model’s explanatory power, it might also yield information about the relevance of each dimension in explaining the measured object representation.

One approach would be to explain each measured response channel by a linear combination of the model dimensions. This is known as population or voxel receptive field modeling in the fMRI literature (Dumoulin & Wandell 2008; Kay et al. 2008; Mitchell et al. 2008). It requires estimating one parameter per model dimension for each measured response channel and enables general linear remixing of the model representational space to explain the measured representation. The model representational space can be stretched, squeezed, and sheared along arbitrary dimensions to account for the measured representation. The large number of parameters usually requires the use of strong priors on the weights (implemented, for example, by regularization penalties used in fitting). In the present scenario, for example, fitting over 100 features per model to predict responses to only 96 stimuli, would yield perfect prediction accuracy on the training set due to over fitting. Moreover, the fit would not be unique without a prior on the weights. Here we take the alternative approach of weighted representational modeling (Diedrichsen et al. 2011), where a single weight is fitted for each model dimension. In this approach, the model representational space can be stretched and squeezed along its original dimensions. However, it cannot be stretched or squeezed along oblique dimensions or sheared. Weighted representational modeling has the advantage of giving more stable and interpretable fits and being directly applicable to similarity judgments. Importantly, it does not require a prior on the weights (i.e. no regularization penalty), which would bias the estimated weights. Our particular approach to weighted representational modeling follows Khaligh-Razavi et al. (2014), using non-negative least squares and cross-validation across images.

Imagine we had an RDM based on spike counts from a population of neurons. If we found the weights by which to multiply the values on each dimension, so as to optimally predict the neuronal data RDM, we would have an indication of the variance each dimension explains in the representational space (resulting from the number of neurons responding to that dimension and the gain of the neuronal responses with respect to that dimension).

Because the squared differences simply sum across dimensions in the squared Euclidean distance, weighting the dimensions and computing the RDM is equivalent to a weighted sum of the single-dimension RDMs. When a dimension is multiplied by weight w, then the squared differences along that dimension are multiplied by w^2^. We can therefore perform the fitting on the RDMs, finding the non-negatively weighted average of the single-dimension model RDMs that best explains the RDM of the measured representation (Figure 1; Khaligh-Razavi et al. 2014). Equation (1) shows that the weights for the model dimensions in the original space can be obtained by taking the square root of the non-negative weights that are estimated for the single-dimension model RDMs.

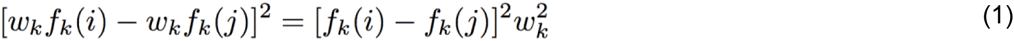

Where is the weight given to dimension, *k* is the value on dimension for stimulus, and is the value on dimension for stimulus. In our case, values are either 0 (absent) or 1 (present). We usedsquared Euclidean distances as the representational dissimilarity measure. The brain RDMs were computed using correlation distance, which is equivalent to the squared Euclidean distance computed for normalized representational patterns.

We estimated the single-dimension model RDM weights with a non-negative-least-squares fitting algorithm (Lawson & Hanson, 1974; also see Khaligh-Razavi & Kriegeskorte, 2014) in Matlab (function lsqnonneg).In order to prevent positive bias of the model performance estimates due to over fitting to a particular set of images, model predictions accuracy was estimated by cross-validation with a subset of the images held out on each fold. For each cross validation fold, we randomly selected 88 of the 96 images as the training set, and used the corresponding pair wise dissimilarities for estimating the model weights. The model weights were then used to predict the pair wise dissimilarities for the eight left-out images. This procedure was repeated until predictions were obtained for all pair wise dissimilarities.

### 2.5 Comparing the explanatory power of categorical and feature-based models

#### 2.5.1 Visualization of the model predictions

To get an impression of the stimulus information that the fitted models can represent, we show the model predictions in Figures 6, 8, and 10. Model predictions are shown for the category model, the feature-based model, and a combined model, which contains all 234 category and feature-based dimensions. The figures also show the data RDMs (IT, EVC, and similarity judgments, respectively) that the models were fitted to, as well as the residual dissimilarity variance that cannot be explained by the models. The residuals were computed by subtracting the predicted dissimilarities from the data dissimilarities. Before subtracting, the predicted and data RDM were each separately rank-transformed and scaled into [0 1], so that the residuals lie in the range [-1 1], or [-100 100] if expressed in dissimilarity percentiles.

**Figure 6.**
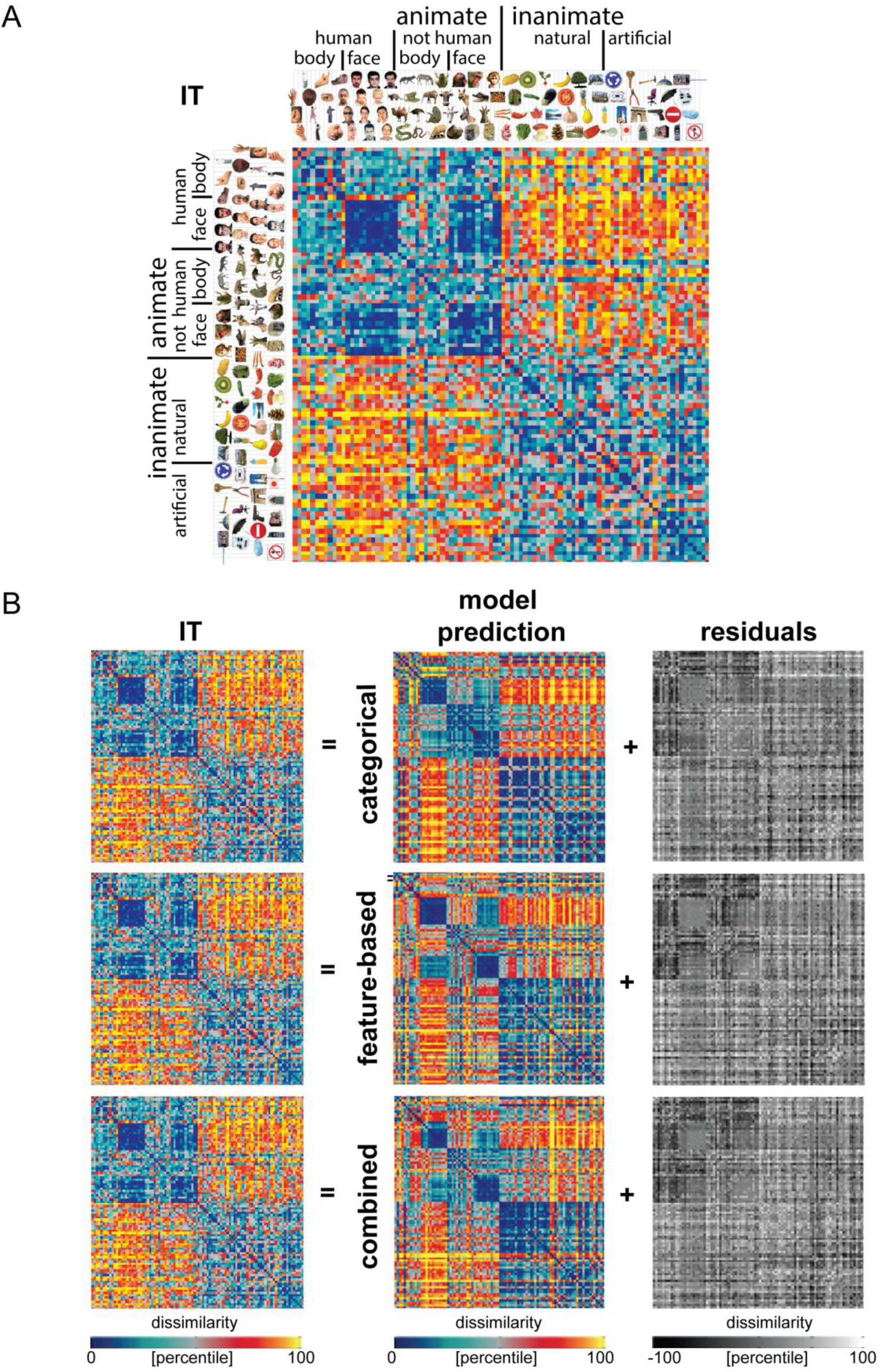
I Model predictions of the IT object representation. A The IT RDM shows a prominent animate/inanimate division, and a face cluster within the animates. The RDM is based on fMRI data from 4 human subjects, averaged at the level of the dissimilarities. Each entry of the RDM represents IT activity-pattern dissimilarity (1 – Pearson’s r; 316 most visually-responsive bilateral IT voxels defined using independent data). The RDM was transformed into percentiles for visualization (see color bar). **B** Model predictions of the IT representation, after weighting the single-dimension model RDMs to optimally predict the IT representation (using independent data). Data and model-prediction RDMs were transformed into percentiles for visualization (see color bar). The residuals were computed based on the transformed RDMs, and highlight which components of the IT RDM could not be explained by the models.

**Figure 7.**
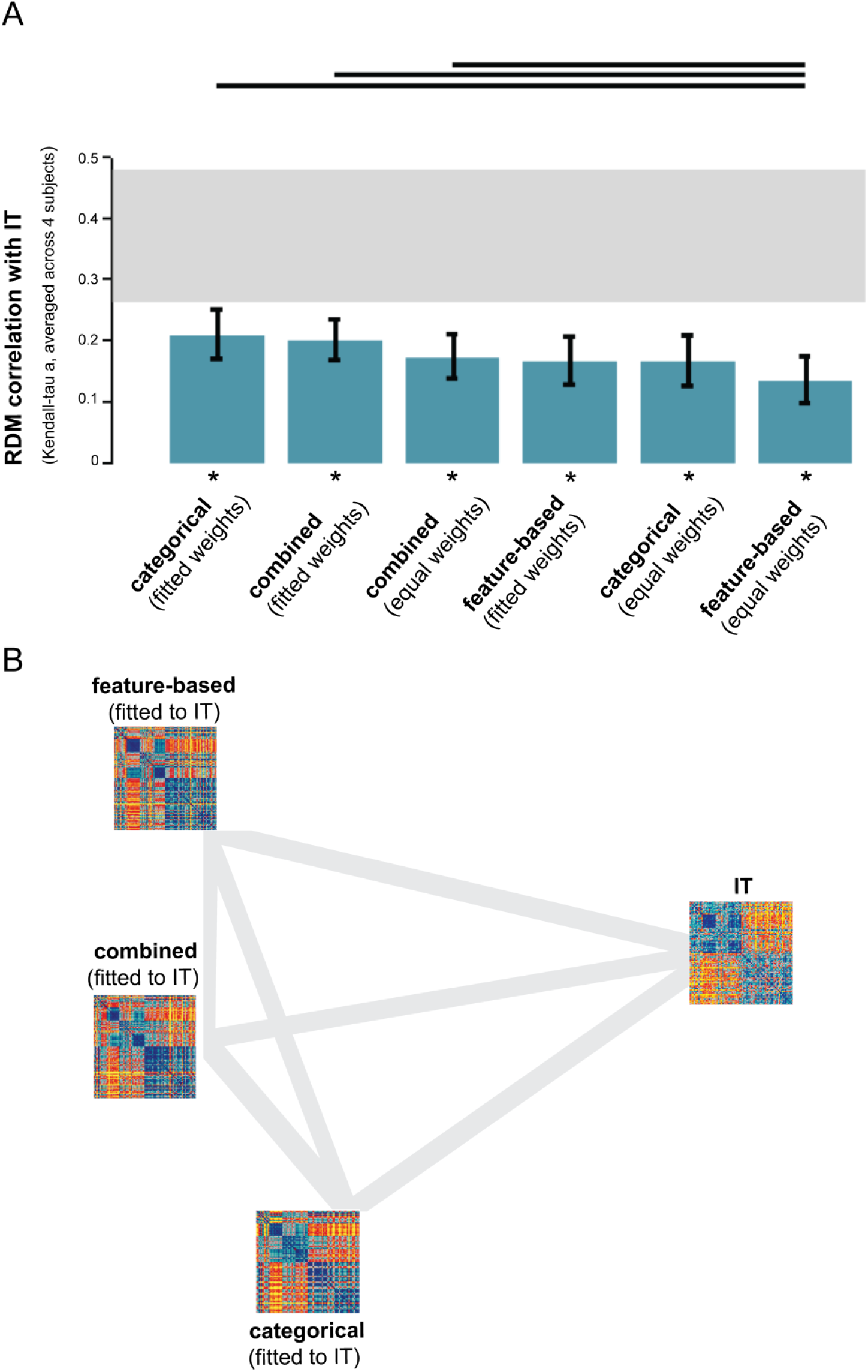
Model performance for IT: the categorical and feature-based models perform equally well. A The bar graphs show the correlation between the IT RDM and each of the model RDMs. Significant correlations between a model RDM and the IT RDM are indicated by an asterisk (stimulus-label randomization test, p <0.05 corrected). Significant differences between models in how well they can account for the IT representation are indicated by horizontal lines plotted above the bars (stimulus-bootstrap test, p < 0.05 corrected). Error bars show the standard error of the mean based on the bootstrap resampling of the stimulus set. The thick gray bar represents the noise ceiling. **B** The multidimensional scaling plot (criterion: metric stress; distance measure: 1-r, where r is Spearman correlation coefficient) visualizes the relationships between the IT RDM and the RDMs predicted by the fitted models. Distances between RDMs reflect their dissimilarity. The thickness of the lines reflects the inevitable distortions that are introduced by dimensionality reduction.

**Figure 8.**
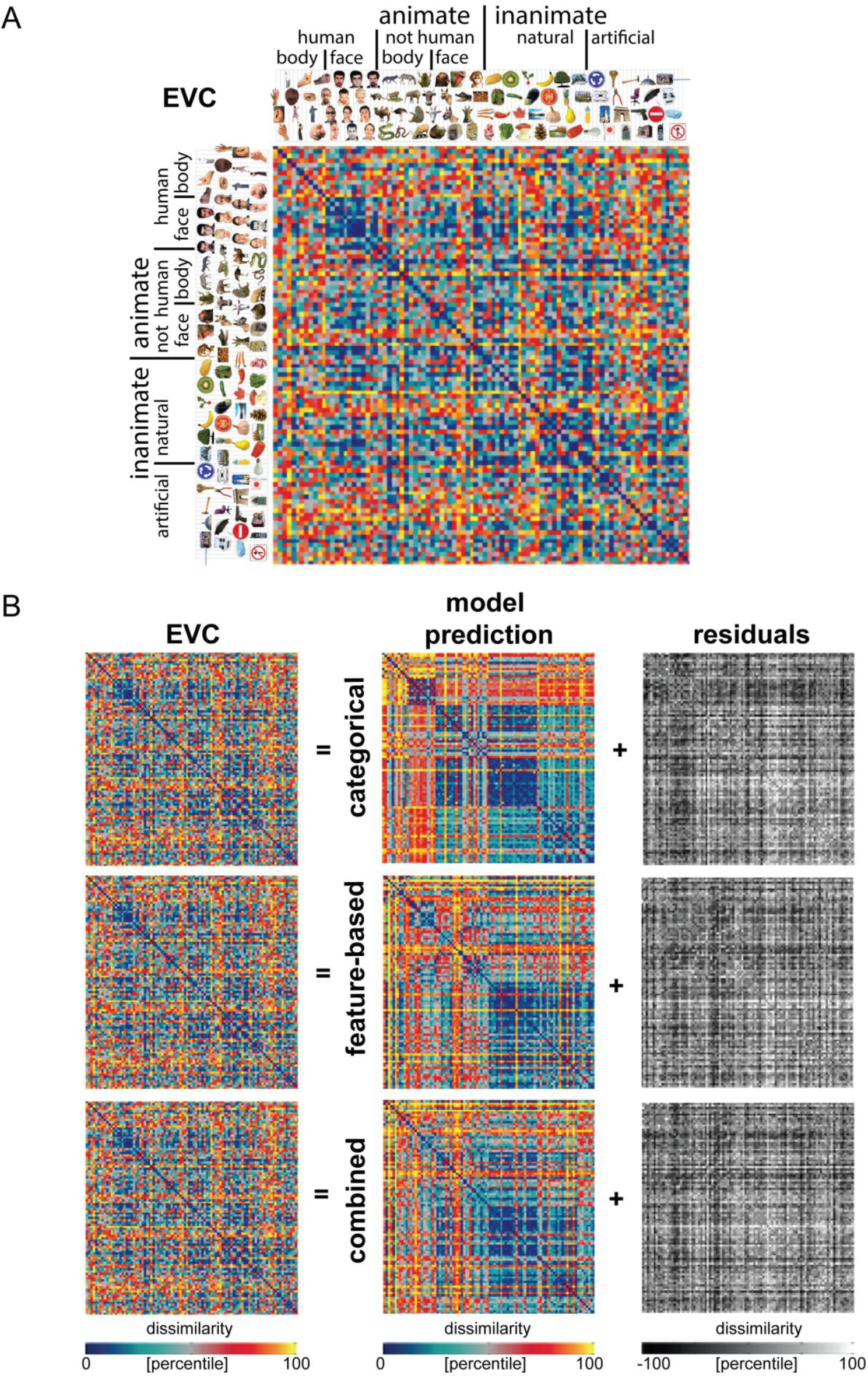
Model predictions of the EVC object representation. A The EVC RDM does not show a clear categorical structure, except for a very weak cluster of human faces. The RDM is based on fMRI data from 4 human subjects, averaged at the level of the dissimilarities. Each entry of the RDM represents EVC activity-pattern dissimilarity (1 – Pearson’s r; 1057 most visually-responsive bilateral EVC voxels defined using independent data). The RDM was transformed into percentiles for visualization (see color bar). **B** Model predictions of the EVC representation, after weighting the single-dimension model RDMs to optimally predict the EVC representation (using independent data). Data and model-prediction RDMs were transformed into percentiles for visualization (see color bar). The residuals were computed based on the transformed RDMs.

**Figure 9.**
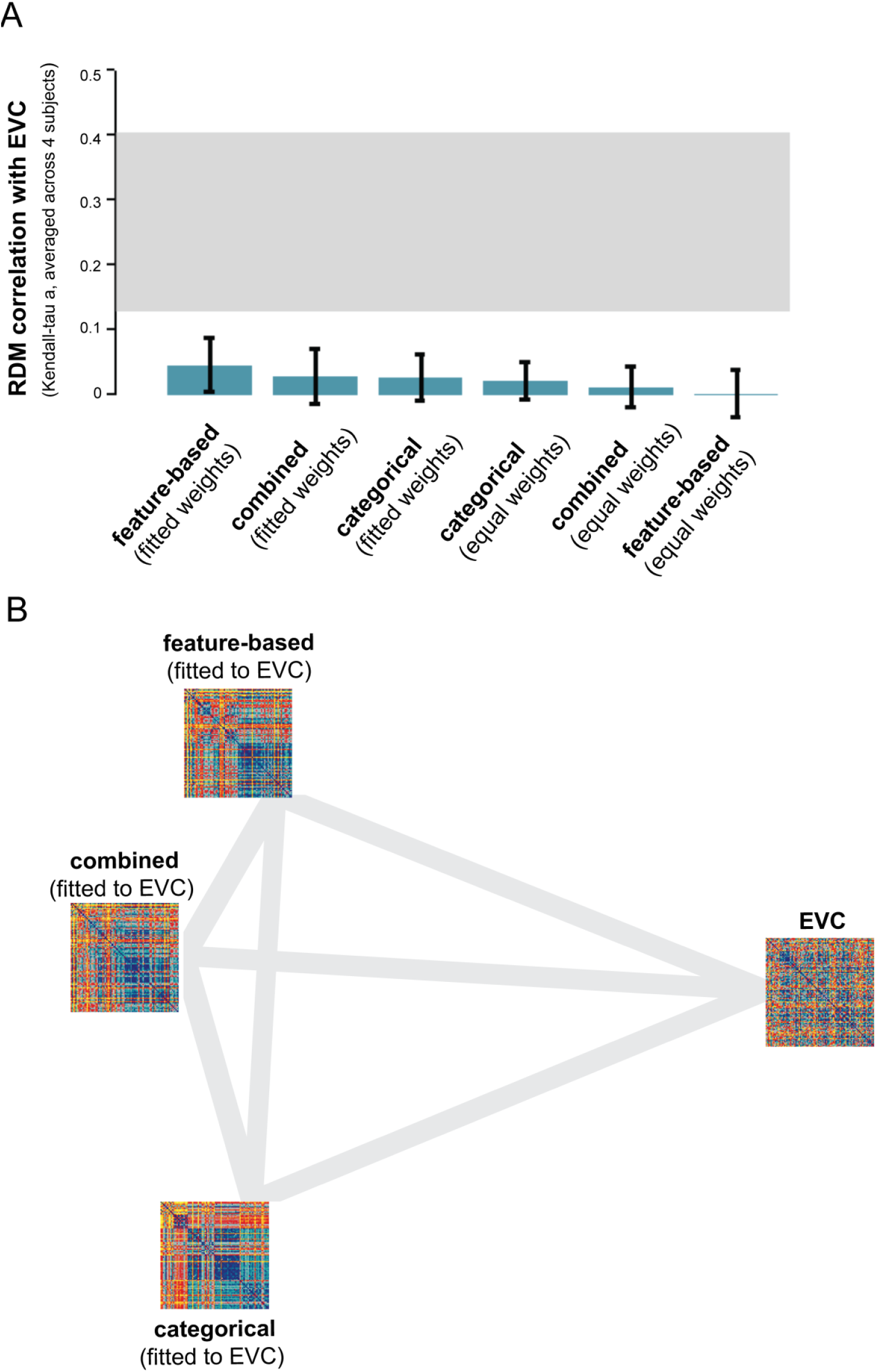
Model performance for EVC: none of the models can explain the EVC representation. A The bar graphs show the correlation between the EVC RDM and each of the model RDMs. Significant correlations between a model RDM and the EVC RDM are indicated by an asterisk (stimulus-label randomization test, p <0.05 corrected). Significant differences between models in how well they can account for the EVC representation are indicated by horizontal lines plotted above the bars (stimulus-bootstrap test, p < 0.05 corrected). Error bars show the standard error of the mean based on the bootstrap resampling of the stimulus set. The gray bar represents the noise ceiling, which indicates the expected performance of the true model given the noise in the data. **B** The multidimensional scaling plot (criterion: metric stress; distance measure: 1-r, where r is Spearman correlation coefficient) visualizes the relationships between the EVC RDMand the RDMs predicted by the fitted models. Distances between RDMs reflect their dissimilarity. The thickness of the lines reflects the inevitable distortions that are introduced by dimensionality reduction.

**Figure 10.**
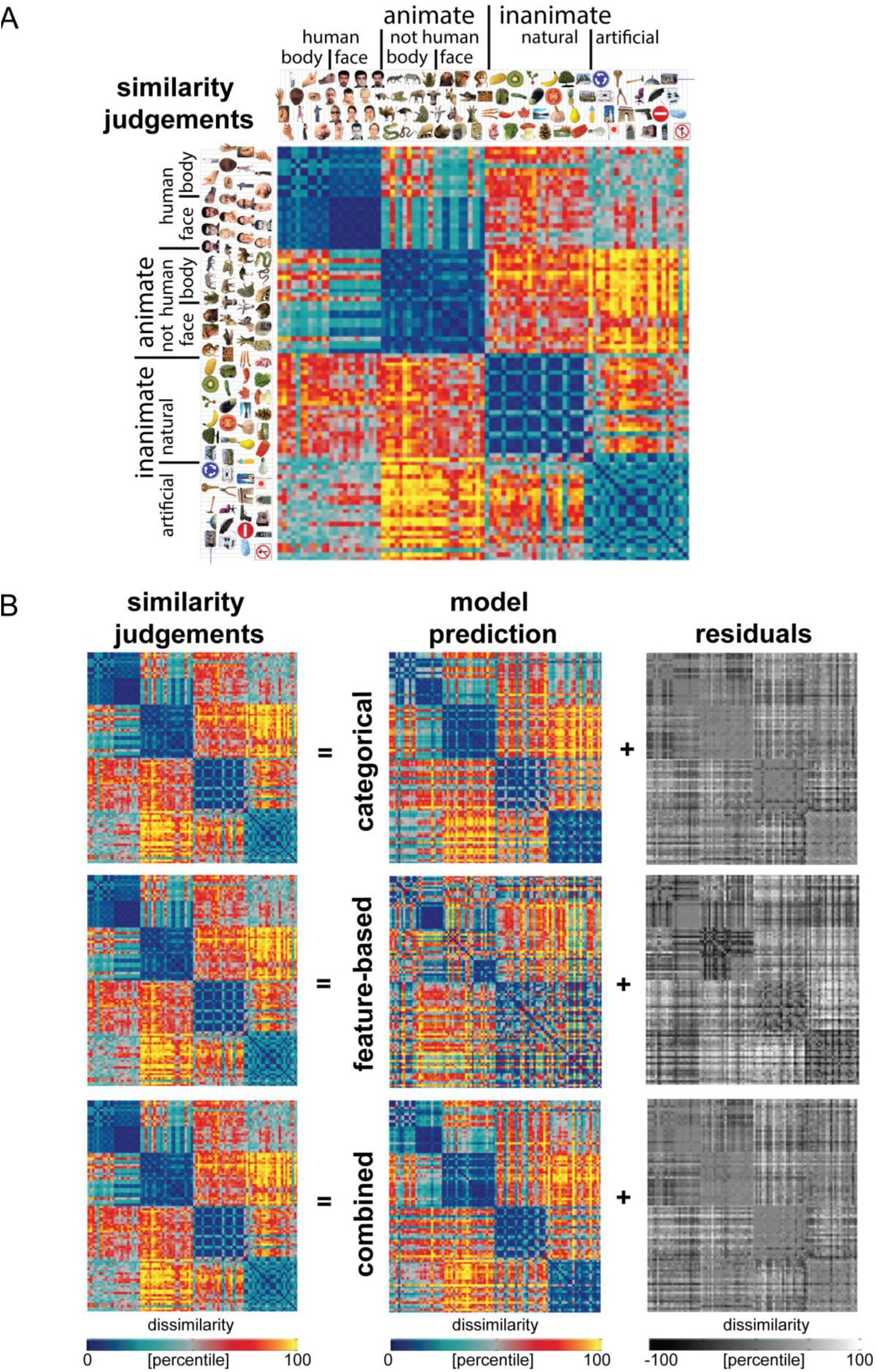
Model predictions of the similarity judgments. A The similarity-judgment RDM shows four main clusters corresponding to humans, non-human animals, natural objects, and manmade objects, and a tight cluster of human faces. The RDM is based on similarity judgments from 16 human subjects, averaged at the level of the dissimilarities. Each entry of the RDM represents the judged dissimilarity between two images. The RDM was transformed into percentiles for visualization (see color bar). **B** Model predictions of the similarity judgments, after weighting the single-dimension model RDMs to optimally predict the similarity judgments (using independent data). Data and model-prediction RDMs were transformed into percentiles for visualization (see color bar). The residuals were computed based on the transformed RDMs.

#### 2.5.2 Inferential analysis on model performance

We used the representational similarity analysis (RSA) toolbox for inferential analyses (Nili et al. 2014). We quantified model performance by measuring the correlation between the data dissimilarities and the dissimilarities predicted by the models. We used Kendall’s rank correlation coefficient tau a as the correlation measure. For each model, we computed the correlation coefficient between each subject’s data RDM and the RDM predicted by the model. Panels A of Figures 7, 9, and 11 show the subject-average correlation coefficients for the fitted (“fitted weights”) as well as the non-fitted (“equal weights”) models.

**Figure 11.**
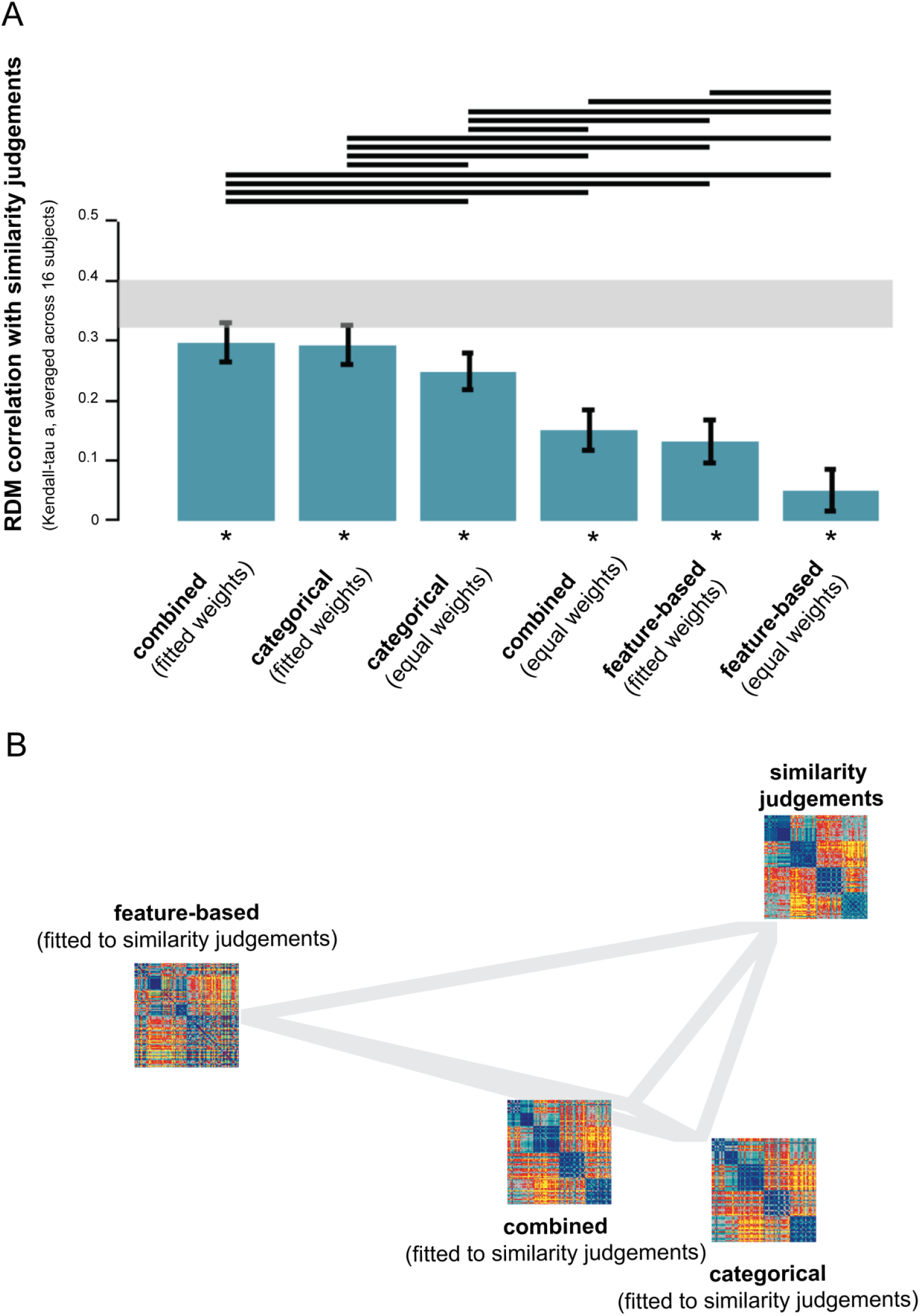
Model performance for similarity judgments: the categorical model outperforms the feature-based model. A The bar graphs show the correlation between the similarity-judgment RDM and each of the model-prediction RDMs. Significant correlations between a model-prediction RDM and the similarity-judgment RDM are indicated by an asterisk (stimulus-label randomization test, p <0.05 corrected). Significant differences between models in how well they can account for the similarity judgments are indicated by horizontal lines plotted above the bars (stimulus-bootstrap test, p < 0.05 corrected). Error bars show the standard error of the mean based on the bootstrap resampling of the stimulus set. The thick gray bar represents the noise ceiling. **B** The multidimensional scaling plot (criterion: metric stress; distance measure: 1-r, where r is Spearman correlation coefficient) visualizes the relationships between the similarity-judgment RDM and the RDMs predicted by the fitted models. Distances between RDMs reflect their dissimilarity. The thickness of the lines reflects the inevitable distortions that are introduced by dimensionality reduction.

We first determined whether each of the model-prediction RDMs is significantly related to each subject-average data RDM using a stimulus-label randomization test (10,000 randomizations per test). This test simulates the null hypothesis of unrelated RDMs (zero correlation). If the actual correlation falls within the top tail of the simulated null distribution, we conclude that the model-prediction and data RDM are significantly related. We corrected for multiple (six) comparisons by controlling the expected false discovery rate at 0.05. We subsequently tested for differences in model performance. We performed pair wise model comparisons using bootstrap re sampling of the stimulus set (1,000 bootstrap re samplings per test). This simulates the variability of model performance across random samples of stimuli. If zero lies in the tail of the simulated distribution of model-performance differences, we conclude that the actual model performances significantly differ from each other. In other words, we conclude that one model can explain the data better than the other. We corrected for multiple (fifteen) comparisons by controlling the expected false discovery rate at 0.05.

The relationships between the data RDMs and the RDMs predicted by the fitted models are visualized in panels B of Figures 7, 9 and 11. The RDMs reside in a high-dimensional space, spanned by the number of dissimilarities contained in the RDM. The distances between RDMs in this space are indicative of their relatedness, i.e. similar RDMs will be placed close together. Because a high-dimensional space is difficult to visualize, we used multidimensional scaling (MDS; criterion: metric stress; distance measure: 1-r, where r is Spearman correlation coefficient) to place the RDMs in a two-dimensional space which preserves the distances between RDMs as well as possible. The thickness of the gray lines reflects the (minimal) distortions that were introduced by the reduction in dimensionality: thin lines indicate that the actual distance in the high-dimensional space is shorter than displayed; thick lines indicate that the actual distance is longer than displayed.

## 3. Results & Discussion

### 3.1 What dimensions do the categorical and feature-based model consist of?

Figure 2 lists the dimensions of the categorical model, and shows whether they are present or absent for each of the 96 object images. Roughly half of the 114 model dimensions are basic-level categories (Rosch et al. 1976), including “face”, “banana”, and “hammer”. A few model dimensions describe sub-ordinate categories, such as “great dane”. The remaining model dimensions describe super-ordinate categories with increasing levels of abstraction, including “mammal”, “animal”, and “organism/living”. In other words, the model consists of a hierarchically nested set of category labels. Approximately one third of the labels describe merged dimensions. Dimensions were merged when their absent/present profiles across the 96 images were highly correlated (r > 0.9). The merged dimensions consist of semantically similar labels (e.g. “nonliving/manmade”, “boy/child/young”), some of which are expected to be less correlated for larger image sets. On average, each object image was described by 5.1 categorical labels (standard deviation = 2.0).

Figure 3 lists the dimensions of the feature-based model, and shows whether they are present or absent for each of the 96 object images. Roughly two-thirds of the 120 model dimensions are object parts (e.g. “eye”, “arm”, “torso”). The remaining model dimensions describe object shape (e.g. “curved”, “rectangular”), color (e.g. “red”, “green”), and texture (e.g. “stubbly”, “woolly”).Finally, a few of the feature-based dimensions are objects which are part of multi object scenes (e.g. “building”, ”shoes”, “glasses”). These features overlap with some of the basic-level categories listed for the categorical model. However, these overlapping features are only listed as present for the feature-based model if they are part of a multi-object scene. Approximately one fifth of the feature-based labels describe merged dimensions. Dimensions were merged when their absent/present profiles across the 96 images were highly correlated (r > 0.9). The merged dimensions consist of labels describing similar features (e.g. “round/circular”), but also of labels that were each uniquely used to describe a single object (e.g. “purple/seat/wheels” for the office chair). These dimensions are expected to be less correlated for larger image sets. On average, each object image was described by 5.5 feature-based labels (standard deviation = 3.6).

The distinction that we make between feature-based and categorical models roughly maps on to the distinction between part-based and holistic representations. The two distinctions share the idea that IT representations of whole objects must emerge from representations of constituent object parts and features. This idea is supported by evidence which suggests that whole objects might be represented as complex conjunctions of features (e.g. Tsunoda et al. 2001; Erez et al. 2015). The terms “holistic” and “categorical” are related because category membership describes an object at a holistic level. However, a categorical object representation does not only require integration of features into a holistic object, it also requires a certain level of invariance to variations in visual appearance among members of the same category. Both of these requirements might be implemented by distributed population coding in IT (e.g. Tsunoda et al. 2001; Vogels et al. 1999).The relative invariance to within-category variation displayed at the level of IT, as indicated by stepwise response profiles and clustering of activity patterns according to category (e.g. Mur et al. 2012; Kriegeskorte et al. 2008b), has been taken to indicate that the representation is categorical. Our categorical model is inspired by these findings. However, the representation also contains a continuous or non-categorical component, as indicated by graded response profiles and replicable within-category dissimilarity variance (e.g. Mur et al. 2012; Kriegeskorte et al. 2008b). This continuous component hints at an underlying feature-based code, consistent with evidence that IT neurons preferentially respond to visual image features of intermediate complexity (e.g. Tanaka 1996; Yamane et al. 2008).

To enable comparison of the models to the measured object representations, which reflect dissimilarities between objects in brain activity and perception, we computed the dissimilarities between objects along each model dimension. Figures 4 and 5 show the single-dimension model RDMs of the categorical and feature-based model, respectively.

### 3.2 Feature-based and categorical models explain the same component of variance in IT

The IT object representation is shown in Figure 6A. As described previously (Kriegeskorte et al. 2008b), the IT object representation shows a categorical structure, with a top-level division between animate and inanimate objects, and a tight cluster of (human) faces within the animate objects. We fitted three models to the IT representation: the categorical model, the feature-based model, and a combined model which contains all categorical and feature-based single dimension model RDMs. Figure 6B shows the model predictions of the IT representation, as well as the variance unexplained by the models. The categorical model predicts the division between animate and inanimate objects and the cluster of (human) faces within the animate objects. The feature-based model also predicts these two prominent characteristics of the IT representation. The residuals indicate that neither model can fully explain the cluster of animate objects because both models predict relatively high dissimilarities between faces and bodies. This mismatch seems somewhat more pronounced for the feature-based model. The prediction of the combined model looks similar to the prediction of each of the two separate models.

To quantify how well the models explain the IT representation, we correlated the model-prediction RDMs with the IT RDM using Kendall’s tau a. We included both the fitted models (“fitted weights”) and the non-fitted models (“equal weights”). We used a stimulus-label randomization test to determine for each model whether its prediction was significantly correlated to the IT RDM. Figure 7A shows that each of the model-prediction RDMs is significantly related to the IT RDM. However, none of the models reaches the noise ceiling, suggesting that the models can still be improved. The noise ceiling indicates the expected performance of the true model given the noise in the data (Nili et al. 2014). We subsequently tested which models performed better than others using bootstrap resampling of the stimulus set. The pair wise model comparisons show that the non-fitted feature-based model performs worse than several other models, namely the fitted categorical model and the fitted and non fitted combined model. No other model comparisons are significant. Importantly, this indicates that the fitted feature-based and fitted categorical model perform equally well. Furthermore, among the fitted models, combining the two models does not improve model performance. This suggests that the feature-based and categorical models explain overlapping variance in the IT object representation. This is consistent with the observation that the two models generate similar predictions (Figure 6B). The multidimensional scaling (MDS) plot shown in Figure 7B further supports the results. The MDS plot visualizes the relationships between the fitted-model predictions and the IT representation. Distances between the representations reflect dissimilarity, such that similar representations are placed close together and dissimilar representations are placed further apart. The three models are approximately equally far away from the IT representation.

We previously showed that objects that elicit similar activity patterns in IT tend to be judged as similar by humans (Mur et al. 2013). This suggests that the IT representation might be predicted from perceived object similarity. Can object-similarity judgments explain the IT representation equally well as the feature-based and categorical models? We repeated our analysis, this time including the similarity judgments as a model. The model “dimensions” in this case are individual subjects (16 in total). Results are shown in Supplementary Figure 2. The pair wise model comparisons show that the similarity judgments can explain the IT representation equally well as the fitted feature-based and fitted categorical models. The fitted similarity judgments perform better than several other models, namely the non-fitted feature-based model, the non-fitted categorical model, and the non-fitted similarity judgments. The finding that the fitted similarity judgments outperform the non-fitted similarity judgments indicates that fitting significantly improves the prediction.

We performed the same analysis for early visual cortex (EVC), which serves as a control region. The EVC representation does not show a strong categorical structure, except for a very weak cluster of human faces (Figure 8A).After fitting the models to the EVC representation, the categorical model predicts a weak cluster of human faces, but none of the models seem to be able to adequately predict the EVC representation (Figure 8B). This observation is confirmed by inferential analyses. Figure 9 shows that none of the model-prediction RDMs is significantly related to the EVC RDM. In other words, none of the models can explain the EVC representation. We repeated this analysis, including the similarity judgments as a model. Supplementary Figure 3 shows that the similarity judgments also cannot explain the EVC representation. This suggests that the feature-based and categorical models, as well as the similarity judgments, capture stimulus information that is not emphasized at the level of EVC. This is consistent with EVC’s known functional selectivity for lower-level image properties such as oriented lines and edges (Hubel & Wiesel 1968).

### 3.3 The categorical model almost fully explains similarity judgments, outperforming the feature-based model

The object-similarity judgments are shown in Figure 10A. As described previously (Mur et al. 2013), the similarity judgments show a categorical structure that reflects and transcends the IT object representation. The judgments reflect the division between animate and inanimate objects that is prominent in the IT representation, and also show a tight cluster of human faces. However, in addition, the similarity judgments emphasize human-related category divisions, including the division between human and non-human animals, and between manmade and natural objects. Figure 10B shows the model predictions of the similarity judgments, and the residual variance unexplained by the models. The prediction of the categorical model shows a close match to the similarity judgments, with four main clusters corresponding to humans, nonhuman animals, natural objects, and manmade objects, and a tight cluster of human faces. The feature-based model cannot predict the four main category clusters prevalent in the similarity judgments, but it can predict the division between animate and inanimate objects and the tight clusters of human and animal faces, which the similarity judgments share with the IT representation.

As shown in Figure 11A, each of the model-prediction RDMs is significantly related to the similarity judgments. Performance of the fitted categorical and combined models approaches the noise ceiling, suggesting that these models can almost fully explain the similarity judgments. The pair wise model comparisons show that these two models outperform all other models, including the fitted feature-based model. This finding suggests that the categorical model can explain variance in the similarity judgments that the feature-based model cannot explain. This is consistent with the observation that the feature-based model cannot predict the four main category clusters prevalent in the similarity judgments. The next best model is the non-fitted categorical model, followed by the non-fitted combined model and the fitted feature-based model. The latter two models each outperform the non-fitted feature-based model, which is ranked last. The fact that each fitted model outperforms its non-fitted counterpart suggests that fitting significantly improves the prediction. The MDS plot in Figure 11B further supports the results, showing that the categorical and combined model are more closely related to the similarity judgments than the feature-based model.

We previously showed that objects that elicit similar activity patterns in IT tend to be judged as similar by humans (Mur et al. 2013). In other words, perceived object similarity can be predicted from the IT object representation. How does the explanatory power of the IT representation compare to that of the categorical and feature-based models? We repeated our analysis, this time including the IT representation as a model. The model “dimensions” in this case are individual subjects (4 in total). Results are shown in Supplementary Figure 4. The pair wise model comparisons show that the IT representation can explain the similarity judgments equally well as the fitted feature-based model. However, the fitted categorical and combined models outperform the IT representation in explaining the similarity judgments. This finding is consistent with the observation that the similarity judgments emphasize several human-related category divisions that can be predicted by the categorical model but that are not present in the IT representation. In sum, our findings suggest that certain aspects of the stimulus information emphasized by the similarity judgments cannot be captured by visual features.

The fact that the performance of the categorical model approaches the noise ceiling indicates that there is not much room for model improvement. This is consistent with the observation that the categorical model falls within the range of inter-subject variability of the similarity judgments (Supplementary Figure 5C). In other words, the single-subject similarity judgments do not seem more similar to each other than to the categorical model. For EVC, and to a lesser extent for IT, this is not the case: the models appear further away from the single-subject data (Supplementary Figure 5A and B). This suggests that the models can still be improved, and corroborates the fact that model performance does not reach the noise ceiling for EVC or IT.

### 3.4 Visual features as stepping stones toward semantics

We found that features, categories, and the combined model explained about equal (and not significantly different) amounts of IT representational variance. The fact that the feature-based model did not explain significant additional variance when added to the categorical model, and vice versa, implies that the two models share the variance that they explain. The explanatory power of both models thus derives from their shared variance component (see Figure 12). This is consistent with the idea that visual features correlated with categorical divisions account for the IT representation, whereas features unrelated to categories do not. This idea is in line with earlier proposals that IT contains feature detectors optimized for category discrimination (Sigala & Logothetis 2002; Ullman et al. 2002; Ullman 2007; Lerner et al. 2008; Rice et al. 2014). Whereas previous studies have focused on disentangling the contribution of features and categories to the IT representation (e.g. Baldassi et al. 2013), our results unite the two by suggesting that the visual features represented in IT might serve as stepping stones toward a representation that emphasizes categorical boundaries or higher-level semantic dimensions.

**Figure 12.**
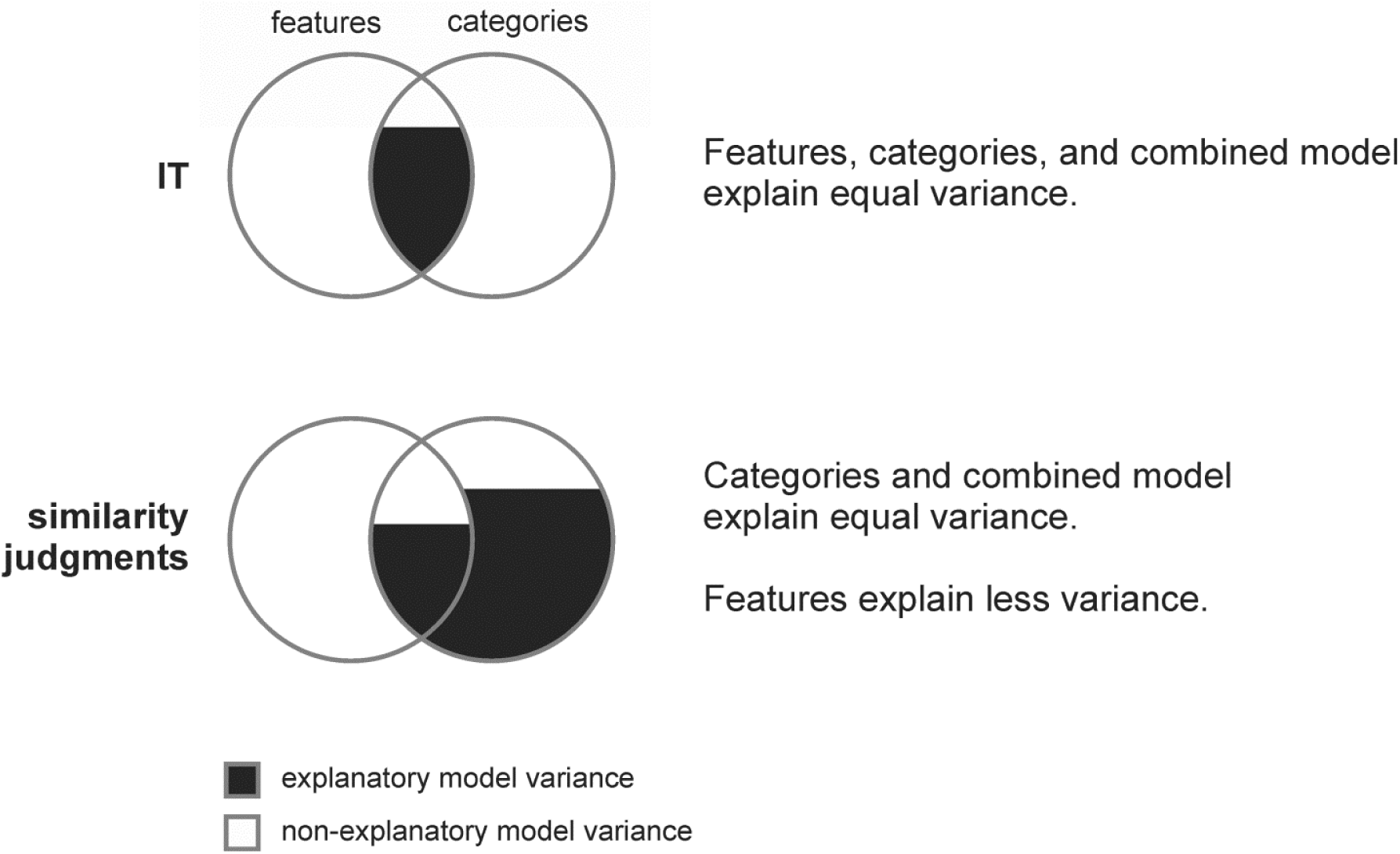
Features correlated with categories explain the IT representation and judgments reflect additional categorical variance. We found that features, categories, and the combined model explained about equal (and not significantly different) amounts of IT representational variance. This implies that the category model does not explain additional variance not explained by the feature model and vice versa. The explanatory power of both models thus derives from their shared variance component. This is consistent with the idea that visual features correlated with categorical divisions account for the IT representation, whereas features unrelated to categories do not. For similarity judgments, the categorical model explained most of the variance and the feature model explained significant, but significantly less variance. The feature model did not explain significant additional variance when added to the category model, implying that the variance it explains is shared with the category model.

For the similarity judgments, the categorical model explained most of the variance and the feature-based model explained significant, but significantly less variance. Furthermore, the feature-based model did not explain significant additional variance when added to the categorical model, implying that the variance it explains is shared with the categorical model (see Figure 12). These findings suggest that the similarity judgments contain categorical variance that is not explained by visual features, reflecting a higher-level more purely semantic representation. Our results further elucidate the nature of the previously reported relationship between the IT object representation and the similarity judgments (Mur et al. 2013). Specifically, they suggest that the dissimilarity variance that each can explain in the other must be driven by the shared variance component of features and categories.

### 3.5 Which model dimensions contribute most to explaining the object representations?

Fitting the models to the measured object representations not only increases the models’ explanatory power, it might also yield information about the relevance of each dimension in explaining the measured object representation, as indicated by the weight that each dimension receives during fitting. In the ideal scenario of spike count measurements for an infinite set of images, the weights would give an indication of the variance each dimension explains in the representational space (resulting from the number of neurons responding to that dimension and the gain of the neuronal responses with respect to that dimension).In the current study, we are several steps away from this ideal scenario. First, we analyze fMRI data. fMRI voxels might not sample the dimensions of the underlying neuronal representational space equally (Kriegeskorte et al. 2010). This compromises the interpretability of the weights. Second, the number of images was limited to 96. This increases multicollinearity between the model predictors. Multicollinearity does not reduce model performance, however, it decreases the stability of the weights. In addition, due to the limited number of images, many dimensions only applied to one particular image. It is unclear to what extent the weights that these dimensions receive during fitting generalize to new images.

Given these considerations, we performed an exploratory analysis on the dimension weights. We first determined, for each of the measured object representations, which of the single dimension model RDMs were significantly related to the representation. This gives an indication of the relevance of the dimensions in explaining the representation when each dimension is considered in isolation. We computed Kendall’s rank correlation coefficient tau a between each single-dimension model RDM and the data RDM, and performed inference by bootstrap resampling the stimulus set (1,000 resamplings, p <0.05 corrected). Supplementary Figure 6 displays the categories and features whose model RDMs show a significant correlation with the IT representation, and with the similarity judgments, respectively. The font size of the category and feature-based labels reflects the relative strength of their correlation with the data dissimilarities. For both the IT representation and the similarity judgments, relevant category labels include super-ordinate categories such as “organism/living”, “nonliving/manmade”, “animal”, “face”, and “food/edible”. The feature-based label “head” is prominently present for both the IT representation and the similarity judgments. Further relevant feature-based labels include labels correlated with animacy or the presence of a face for the IT representation (e.g. “skin”, “hair”, “nose/mouth”) and labels describing object shape and color for the similarity judgments (e.g. “symmetrical”, “red”, “green”). Subsequently, we inspected the dimension weights obtained by non-negative least-squares fitting. The dimension weights are shown in Supplementary Figure 7. Only weights for dimensions that applied to more than one image are shown. The 15 to 20 first-ranked dimensions show a reasonable overlap with the dimensions shown in Supplementary Figure 6. These observations are consistent with the idea that IT represents visual features that are informative about category membership. Future studies should use larger image sets and additional inferential procedures to validate the results of our exploratory analysis.

Our results demonstrate the feasibility of weighted representational modeling (Diedrichsen et al. 2011) for fitting models based on image labels obtained from human observers. In weighted representational modeling, a single weight is fitted for each model dimension. In other words, the model representational space can be stretched and squeezed along its original dimensions to best explain the measured representation. This allows less flexibility than population or voxel receptive field modeling (Dumoulin & Wandell 2008; Kay et al. 2008; Mitchell et al. 2008), in which the model representational space can additionally be sheared along arbitrary dimensions to account for the measured representation. However, the increased flexibility of voxel receptive field modeling comes at the cost of a larger number of parameters, i.e. a weight is fitted for each model dimension and each measured response channel. This requires a prior on the weights, which biases the estimated weights. Weighted representational modeling does not require a prior on the weights, and has the advantage of giving more stable and interpretable fits and being directly applicable to similarity judgments.

### 3.6 Conclusion

We have shown that visual features can explain the IT representation to a considerable extent and that categorical predictors do not explain additional IT variance beyond that explained by features. However, only visual features related to categories appeared effective at explaining IT representational variance. This is consistent with IT consisting of visual feature detectors that are designed (by visual development or evolution) to emphasize categorical divisions. Similarity judgments reflect additional categorical variance not explained by visual features. Our results are consistent with the view that IT uses visual features as stepping stones toward a representation that emphasizes categorical boundaries or higher-level semantic dimensions.

We used weighted representational modeling to estimate the contributions of visual features and categories in explaining the IT representation. Weighted representational modeling (Diedrichsen et al. 2011) provides a useful methodology for exploring the degree to which different representational models can explain a representation. Such models have much fewer parameters than voxel/population receptive field models, can be fitted without priors that bias the weight estimates and can be applied directly to representational dissimilarity matrices (including those from human similarity judgments). The particular approach of non-negative least squares with cross-validation across stimuli (Khaligh-Razavi et al. 2014) is shown here to be useful not only for fitting combinations of image-computable model representations, but also for models based on labels obtained from human observers.

## Acknowledgements

We would like to thank SeyedKhaligh-Razavi for sharing his code for the model weighting. This work was funded by the Medical Research Council of the UK (program MC-A060-5PR20), a British Academy Postdoctoral Fellowship to MM, and a Wellcome Trust Project Grant (WT091540MA) and a European Research Council Starting Grant (261352) to NK.

